# Elevated ACE2 expression in the olfactory neuroepithelium: implications for anosmia and upper respiratory SARS-CoV-2 entry and replication

**DOI:** 10.1101/2020.05.08.084996

**Authors:** Mengfei Chen, Wenjuan Shen, Nicholas R. Rowan, Heather Kulaga, Alexander Hillel, Murugappan Ramanathan, Andrew P. Lane

## Abstract

The site of SARS-CoV-2 entry and replication critically impacts strategies for COVID-19 diagnosis, transmission mitigation, and treatment. We determined the cellular location of the SARS-CoV-2 target receptor protein, ACE2, in the human upper airway, finding striking enrichment (200-700 folds) in the olfactory neuroepithelium relative to nasal respiratory or tracheal epithelial cells. This cellular tropism of SARS-CoV-2 may underlie its high transmissibility and association with olfactory dysfunction, while suggesting a viral reservoir potentially amenable to intranasal therapy.

## Introduction

The ongoing outbreak of coronavirus disease 2019 (COVID-19), caused by the severe acute respiratory syndrome coronavirus 2 (SARS-CoV-2), has become a major threat to global health [1]. The mechanism of cellular entry by SARS-CoV-2 is through binding to angiotensin-converting enzyme 2 (ACE2) [2, 3], a metalloproteinase ectoenzyme that primarily functions in the regulation of angiotensin II, but also has non-catalytic roles such as intestinal neutral amino acid transport. The level of ACE2 protein and its subcellular localization in the respiratory tract may be a key determinant of susceptibility to infection, symptoms, and outcomes in COVID-19. In humans, ACE2 protein is broadly expressed in the lung, kidney, and small intestine [4]. Pathological analysis of COVID-19 postmortem samples shows substantial damage in the lung [5], suggesting that the airway is the principal entry and target of SARS-CoV-2. However, analysis of multiple single cell RNA-seq datasets reveal overall low ACE2 RNA transcription in nasal airway epithelium, with further reduced expression in lower airway club cells and rare expression in alveolar epithelial type II cells [6]. This pattern of ACE2 expression provides evidence that the upper, rather than the lower, airway is the initial site of SARS-CoV-2 infection.

There is growing interest in a presentation of SARS-CoV-2 infection characterized by olfactory loss without concomitant nasal inflammatory symptoms. Disturbances in the sense of smell have been widely reported in COVID-19 patients internationally, with a reported prevalence as high as 85% in a large, multicenter European survey [7]. These reports show, importantly, that some COVID-19 patients manifest olfactory loss as their initial or only symptom. As this presentation is largely not recognized or thought to mandate isolation, this patient group may be a source of continued viral spread and a target population for early intervention and mitigation. The loss of the sense of smell suggests the possibility of direct targeting by SARS-CoV-2 of the olfactory system. However, the cellular location of ACE2 protein in the olfactory epithelium has not been previously demonstrated.

## Results

Within the nasal cavity, the specialized olfactory neuroepithelium has an apical surface consisting mainly of sustentacular cells, which support neuronal dendritic projections containing the odor-sensing cilia. We performed immunohistological analysis to determine the location of ACE2 protein in human nasal and trachea biopsies. Confocal images demonstrated that the majority of ACE2 staining is localized to the apical surface of Krt18^+^ sustentacular cells in the olfactory neuroepithelium (Figure 1A and B). This distribution of ACE2 protein is similar to that reported in bat nasal epithelium [8]. We further quantified olfactory ACE2 expression and found the number of ACE2 positive cells to be comparable between healthy controls and chronic rhinosinusitis (CRS), a common inflammatory disease of the nasal mucosa (Figure 1A, C, and H) and affects the olfactory mucosa [9]. ACE2 is not present in olfactory neurons, demonstrated by co-staining with the immature and mature olfactory neuron marker DCX (Figure 1E and F) and PGP9.5 (Figure 1G), respectively.

**Figure 1.**
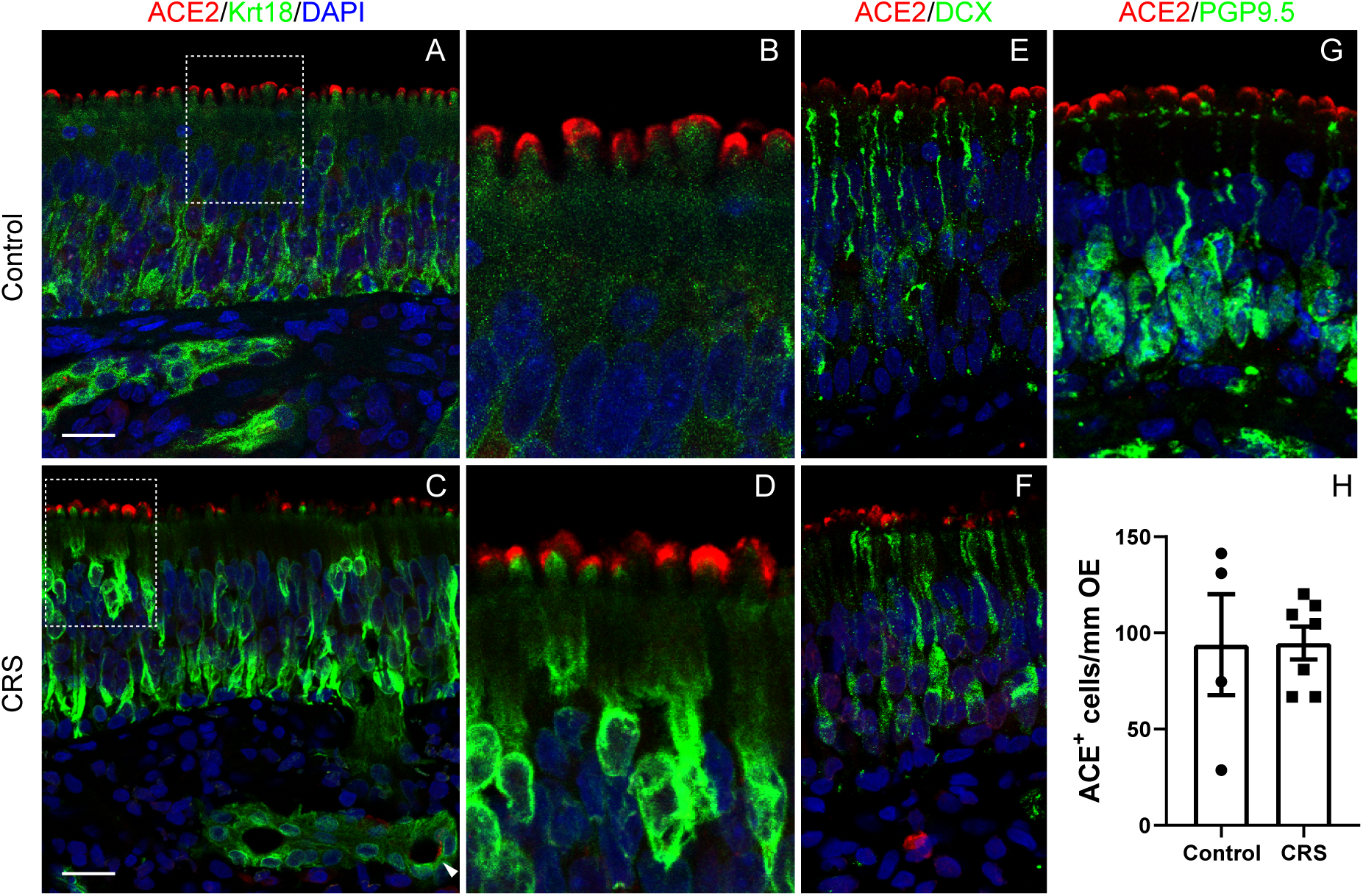
Cellular location of ACE2 in human olfactory epithelium. (A and B) Confocal image of ACE2 (red) and Krt18 (green) immunostaining in the healthy control olfactory neuroepithelium. ACE2 is localized to the apical surface of Krt18 positive sustentacular cells in the olfactory epithelium. (C and D) Representative image of ACE2 and Krt18 immunostaining in the olfactory neuroepithelium of a CRS patient. The Boxed area in Panel A and C was highlighted in B and D, respectively. (E and F) The location of ACE2 and DCX positive immature olfactory sensory neurons in control (E) and CRS biopsy (F). (G) The location of ACE2 and PGP9.5 positive mature olfactory sensory neurons in control. (H) Quantification of ACE2 positive cells in olfactory epithelium. Data are represented as mean ± SEM. *p* value was calculated by unpaired two-tailed Student’s t test (H). Scale bars, 20 µm.

High intensity ACE2 staining was detected in all 11 olfactory mucosal biopsies. In addition, ACE2 is frequently observed in Bowman’s glands (Figure 2A). In the adjacent nasal respiratory epithelium, ACE2 is also located on the apical surface (Figure 2B and C), with a significantly lower level of expression than the olfactory epithelium (Figure 2D). As shown in Figure 2E in the adjacent area, intensive ACE2 expressed in PGP9.5 olfactory region but can barely be detected in PGP9.5^+^ respiratory epithelium. Only 47.4% of nasal respiratory epithelial biopsies (9 in 19) contained ACE2 positive epithelial cells. The intensity of ACE2 fluorescence was 200-700-fold higher in the olfactory epithelium (Figure 2F).

**Figure 2.**
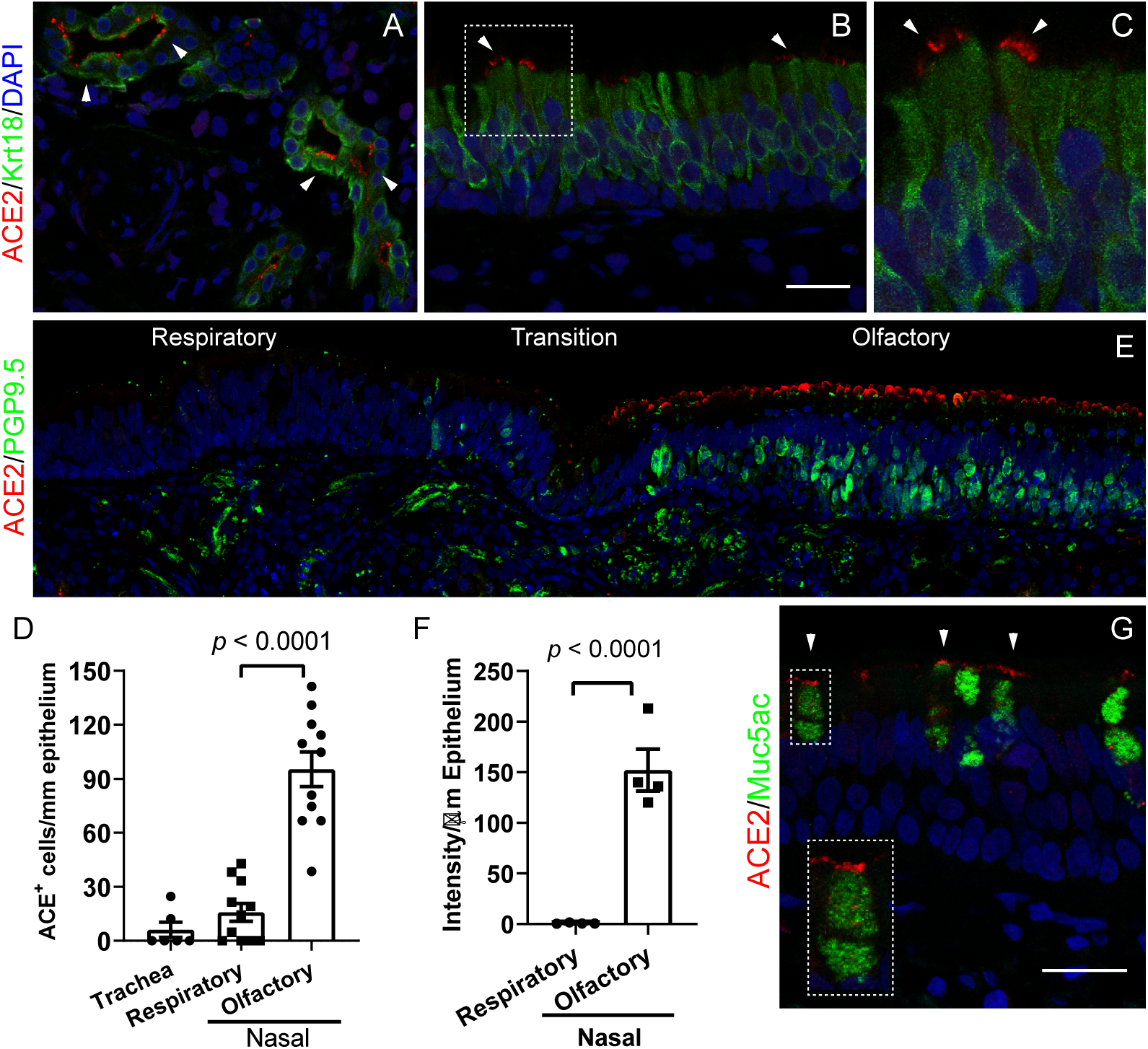
Elevated ACE2 in olfactory relative to respiratory epithelium. (A-C) Expression of ACE2 in glands (A) and nasal respiratory epithelium (B and C). The Boxed area in Panel B was highlighted in C. (D) Quantiication of the ACE2^+^ cell per mm epithelium. The positive cells in each nasal specimen that contained both respiratory and olfactory epithelium were counted. Dots in graph represent independent specimens. Data are represented as mean ± SEM. *p* value was calculated by unpaired two-tailed Student’s t test. (E) Representative image of respiratory-olfactory mucosa adjacent area. PGP9.5 and ACE2 co-staining image was obtained using confocal microscope under the tile scan mode. (F) Quantiication of the ACE2 luorescence intensity per µm epithelium using Image J. (G) Co-staining of ACE2 and secretory cell marker Muc5ac in trachea airway epithelium. The inset represents magniication of the selected area. Scale bars, 20 µm

We further examined ACE2 protein in tracheal epithelium. In 2 of 7 specimens, we detected low expression of ACE2 in Muc5ac^+^ secretory cells (Figure 2G). The expression of ACE2 by secretory cells is reminiscent of the recent findings that increased ACE2 expression in small airway epithelium in COPD patients [10], a disease characterized by secretory cell hyperplasia [11]. Together, the comparatively enhanced human airway expression of ACE2 localized to the olfactory neuroepithelium (Figure 2D and E) suggests a mechanism of olfactory loss and a potential entry point of SARS-CoV-2 into the central nervous system and causes neurological symptoms in COVID-19 patients [12].

## Discussion

Understanding of the pattern of viral load in tissues of COVID-19 patients is critical for diagnosis, management of transmission, and potential treatment strategies. Detection of SARS-CoV-2 in clinical specimens shows that the highest viral copy number is found in nasal swabs (∼200 fold), compared to bronchoalveolar lavage or pharyngeal swabs [13, 14]. In the early stages of SARS-CoV-2 infection, viral RNA can readily detected in upper respiratory specimens but not in blood, urine, or stool [15]. These findings, taken together with ACE2 protein cellular localization presented here, suggests that active virus infection and replication occurs in the apical layer of nasal and olfactory mucosa. The differential expression of ACE2 in the olfactory neuroepithelium and respiratory epithelium may help account for the spectrum of nasal-related symptoms, while also raising the intriguing possibility that COVID-19 may be amenable to novel therapeutic approaches. Whether nasal saline irrigation, a common treatment for sinonasal conditions, is beneficial or counter-productive in SARS-CoV-2 infection remains to be determined; however, consideration should be given to the delivery of topical anti-viral additives, such as detergent or povidone-iodine, directed at the nasal and nasopharyngeal viral reservoirs.

## ONLINE METHODS

### Human Olfactory and Nasal Tissue

The research protocol involving human specimens was approved by the Johns Hopkins institutional review board, and all subjects provided signed informed consent. Human olfactory epithelial or nasal airway epithelial samples were collected from CRS patients and from control subjects undergoing endonasal surgical approaches for non-CRS disease processes. The specimen was immediately fixed in 4% PFA for 5h at 4 °C with gentle rotation. After being washed in PBS, tissues were equilibrated sequentially in 15% and 30% sucrose, the tissue was then embedded in Optimum Cutting Temperature (OCT, Tissue-Tek) for sectioning.

### Immunohistochemistry

After antigen retrieval step, sections were washed in PBS and then blocked in 10% normal serum containing 0.1% Triton X-100, followed by incubation with primary antibodies at 4 °C overnight. After washing in PBS, the tissue sections were incubated with Alexa Fluor conjugated secondary antibodies along with DAPI for nuclear counterstaining. The following primary antibodies were used: Goat anti-ACE2 (1:40, AF933; R&D) Mouse anti-Keratin 18 (1:500, Pierce MA1-39483), Rabbit anti-DCX (1:500, GeneTex, GTX134052), Mouse anti-Mucin 5AC (1:200, Abcam ab3649).

### Confocal Imaging and Quantification

Immunostaining images were obtained using a Zeiss LSM 780 confocal microscope. For quantification, at least 3 images were collected from each specimen using 40x objectives under the z stack scan mode at same depth. Positive cells in olfactory or respiratory mucosa were quantified per mm of surface epithelium. Quantification of the ACE2 fluorescence intensity was preformed using Image J software.

### Statistical analyses

Data are expressed as mean ± SEM. as indicated. Data analyses were carried out using GraphPad Prism. For experiments with two groups, *P* values were calculated using the unpaired two-tailed Student’s t-test. Differences were considered significant when *P* < 0.05.

## ACKNOWLEDGEMENTS

This work was funded by NIH Grants R01 AI132590, R01 DC016106 (A.P.L).

## AUTHOR CONTRIBUTIONS

M.C. and A.P.L. designed research; M.C., W.S., and H.K. collected data, M.C. performed experiment; M.C., W.S., A.H., and A.P.L. analyzed data; M.C., N.R.R., M.R., and A.P.L. wrote the paper.

